# An Eigenvalue Test for spatial Principal Component Analysis

**DOI:** 10.1101/151639

**Authors:** V. Montano, T. Jombart

## Abstract

**Background:** The spatial Principal Component Analysis (sPCA, Jombart 2008) is designed to investigate non-random spatial distributions of genetic variation. Unfortunately, the associated tests used for assessing the existence of spatial patterns (*global and local test*; Jombart et al. 2008) lack statistical power and may fail to reveal existing spatial patterns. Here, we present a non-parametric test for the significance of specific patterns recovered by sPCA.

**Results:** We compared the performance of this new test to the original *global* and *local* tests using datasets simulated under classical population genetic models. Results show that our test outperforms the original *global* and *local* tests, exhibiting improved statistical power while retaining similar, and reliable type I errors. Moreover, by allowing to test various sets of axes, it can be used to guide the selection of retained sPCA components.

**Conclusions:** As such, our test represents a valuable complement to the original analysis, and should prove useful for the investigation of spatial genetic patterns.

## INTRODUCTION

The principal component analysis (PCA; Pearson 1901; Hotelling 1933) is one of the most common multivariate approaches in population genetics (Jombart et al 2009). Although PCA is not explicitly accounting for spatial information, it has often been used for investigating spatial genetic patterns (Novembre and Stephens 2008). As a complement to PCA, the spatial principal component analysis (sPCA; Jombart et al. 2008) has been introduced to explicitly include spatial information in the analysis of genetic variation, and gain more power for investigating spatial genetic structures.

sPCA finds synthetic variables, the principal components (PCs), which maximise both the genetic variance and the spatial autocorrelation as measured by Moran's *I* (Moran 1950). As such, PCs can reveal two types of patterns: ‘*global*' structures, which correspond to positive autocorrelation typically observed in the presence of patches or clines, and ‘*local*’ structures, which correspond to negative autocorrelation, whereby neighboring individuals are more genetically distinct than expected at random (for a more detailed explanation on the meaning of *global* and *local* structures see Jombart et al.. 2008). The *global* and *local* tests have been developed for detecting the presence of global and local patterns, respectively (Jombart et al. 2008). Unfortunately, while these tests have robust type I error, they also typically lack power, and can therefore fail to identify existing spatial genetic patterns (Jombart et al.. 2008). Moreover, they can only be used to diagnose the presence or absence of spatial patterns, and are unable to test the significance of specific structures revealed by sPCA axes.

In this paper, we introduce an alternative statistical test which addresses these issues. This approach relies on computing the cumulative sum of a defined set of sPCA eigenvalues as a test statistic, and uses a Monte-Carlo procedure to generate null distributions of the test statistics and approximate p-values. After describing our approach, we compare its performances to the global and local tests using simulated datasets, investigating several standard spatial population genetics models. Our approach is implemented as the function *spca_randtest* in the package *adegenet* (Jombart 2008; Jombart and Ahmed 2011) for the R software (R Core Team 2017).

## METHODS

### Test statistic

As in most multivariate analyses of genetic markers, our approach analyses a table of centred allele frequencies (i.e. set to a mean frequency of zero), in which rows represent individuals or populations, and columns correspond to alleles of various loci (Jombart et al 2008; Jombart et al 2009; Jombart et al 2010). We note *X* the resulting matrix, and *n* the number of individuals analysed. In addition, the sPCA introduces spatial data in the form of a *n* by *n* matrix of spatial weights *L*, in which the *i*^th^ row contains weights reflecting the spatial proximity of all individuals to individual *i*. The PCs of sPCA are then found by the eigen-analysis of the symmetric matrix (Jombart et al. 2008):

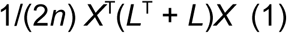

We note *λ* the corresponding non-zero eigenvalues. We differentiate the *r* positive eigenvalues *λ*^*+*^, corresponding to global structures, and the ‘*s*’ negative eigenvalues *λ*^*-*^, corresponding to local structures, so that *λ =* {*λ*^*+*^,*λ*^*-*^}. Without loss of generality, we assume both sets of eigenvalues are ordered by decreasing absolute value, so that 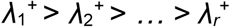 and 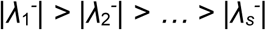. Simply put, each eigenvalue quantifies the magnitude of the spatial genetic patterns in the corresponding PC: larger absolute values indicate stronger global (respectively local) structures. We note 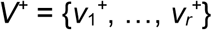 and 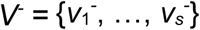 the sets of corresponding PCs.The most natural choice of test statistic to assess whether a given PC contains significant structure would seem to be the corresponding eigenvalue. This would, however, not account for the dependence on previous PCs: 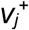(respectively 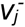) can only be significant if all previous PCs 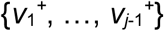 are also significant. To account for this, we define the test statistic for 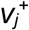 as:

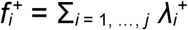

and as:

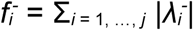

for 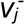.

### Permutation procedure

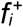 and 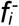 become larger in the presence of strong global or local structures in the first *i*^th^ global / local PCs. Therefore, they can be used as test statistics against the null hypotheses of absence of global or local structures in these PCs. The expected distribution of 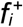 and 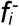 in the absence of spatial structure is not known analytically. Fortunately, it can be approximated using a Monte-Carlo procedure, in which at each permutation individual genotypes are shuffled to be assigned to a different pair of coordinates than in the observed original dataset and 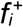 and 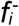 are computed. Note that the original values of the test statistic are also included in these distributions, as the initial spatial configuration is by definition a possible random outcome. The *p*-values are then computed as the relative frequencies of permuted statistics equal to or greater than the initial value of 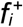 or 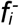.

To guide the selection of global and local PCs to retain, the simulated values of each eigenvalue (from most positive to most negative), which make up the 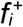 and 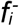 statistics, are also recorded during the permutation procedure. In this way, if global or local structures are detected to be significant, an observed *p*-value for each observed eigenvalue can be estimated by comparison with its simulated eigenvalue distribution. Note that the number of eigenvalues produced by an sPCA does not change between the observed and permutated datasets, so each observed eigenvalue can be compared with the distribution of the corresponding simulated one. This testing procedure can be used with increasing numbers of retained axes. Because each test is conditional on the previous tests, incremental Bonferroni correction is used to avoid the inflation of type I error, so that the significance level for the *i*^th^ PC will be α / i, where α is the target type I error. Hence, the correction implies that if the most positive (or negative) eigenvalue is significant in regards with the chosen *p-*value threshold, the second eigenvalue is tested for a *p-*value threshold that is the half of the previous and so on. The entire testing procedure is implemented in the function *spca_randtest* in the package *adegenet* (Jombart 2008; Jombart and Ahmed 2011) for R (R Core Team 2017). A flow chart of the test procedure is shown in Figure 1.

**Figure 1.**
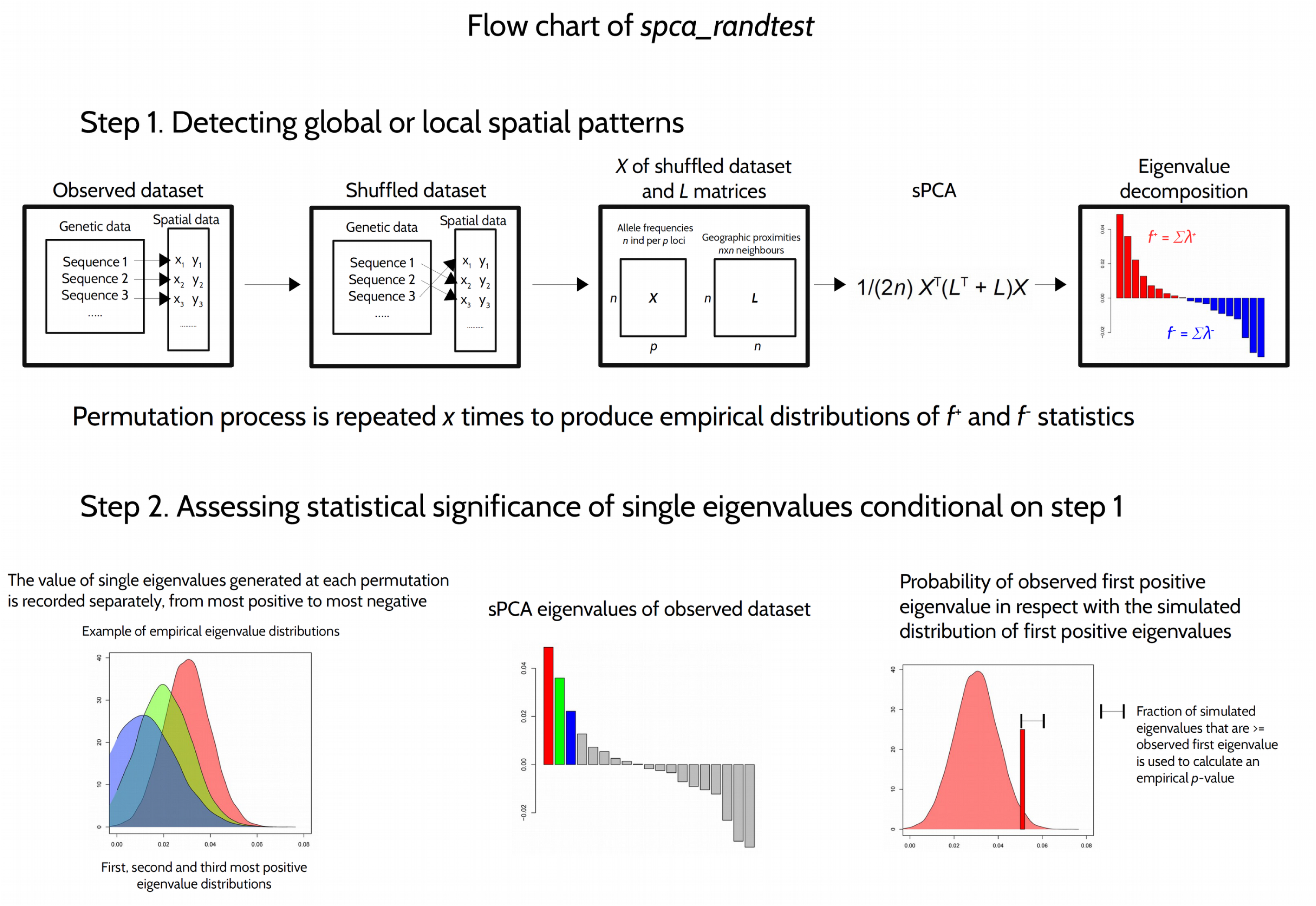
Flow chart illustrating the steps of the *spca_randtest*. The first step on the top panel assess the statistical significance of global either local patterns. If at least one of the two is significant, the second step of the test exploits the eigenvalue distribution recorded over the permutations to obtain an empirical *p*-value for each eigenvalue, starting from the most positive (or most negative). As the first eigenvalue is significant in comparison with a chosen threshold, the following is tested and compared to a more stringent threshold (Bonferroni correction) until a non-significant eigenvalue is found and the routine stops.

### Simulation study

To assess the performance of our test, we simulated genetic data under three migration models: island (IS) and stepping stone (SS), using the software GenomePop 2.7 (Carvajal-Rodríguez 2008), and isolation by distance (IBD), using *IBDSimV2.0* (Leblois 2009). We simulated the IS and SS models with 4 populations, each with 25 individuals, and a single population under IBD with 100 individuals. 200 unlinked biallelic diploid loci (or single nucleotide polymorphisms; SNPs) were simulated. Populations evolved under constant effective population size θ = 20, and interchanged migrants at three different symmetric and homogeneous rates (0.005, 0.01, and 0.1). We performed 100 independent runs for each of the three migration rates, for a total of 300 simulated dataset per migration model.

To quantify type I error rates for the *spca_randtest, global* and *local tests*, we extracted 100 random coordinates from 10 square 2D grids, using the function *spsample* from the *spdep* package (Bivand et al. 2013). In order to evaluate the rate of false negatives for global patterns, we manually generated 10 sets of 100 pairs of coordinates simulating gradients and/or patches from 2D grids. An example of simulated global patterns is presented in Figure 2. To test for the rate of false negatives for local patterns, we perform a principal component analysis on 10 random datasets simulated under the SS model with 0.005 migration rate. We used the coordinates of the individuals on the first principal component and set the second coordinate to zero for all individuals (1D). With the coordinates so produced, we used the function *chooseCN* in adegenet to obtain 10 neighbouring graphs where the most genetically distinct individuals (falling in the upper quartile of the pairwise genetic distances) are considered as neighbors, while the others are non-neighbors.

**Figure 2.**
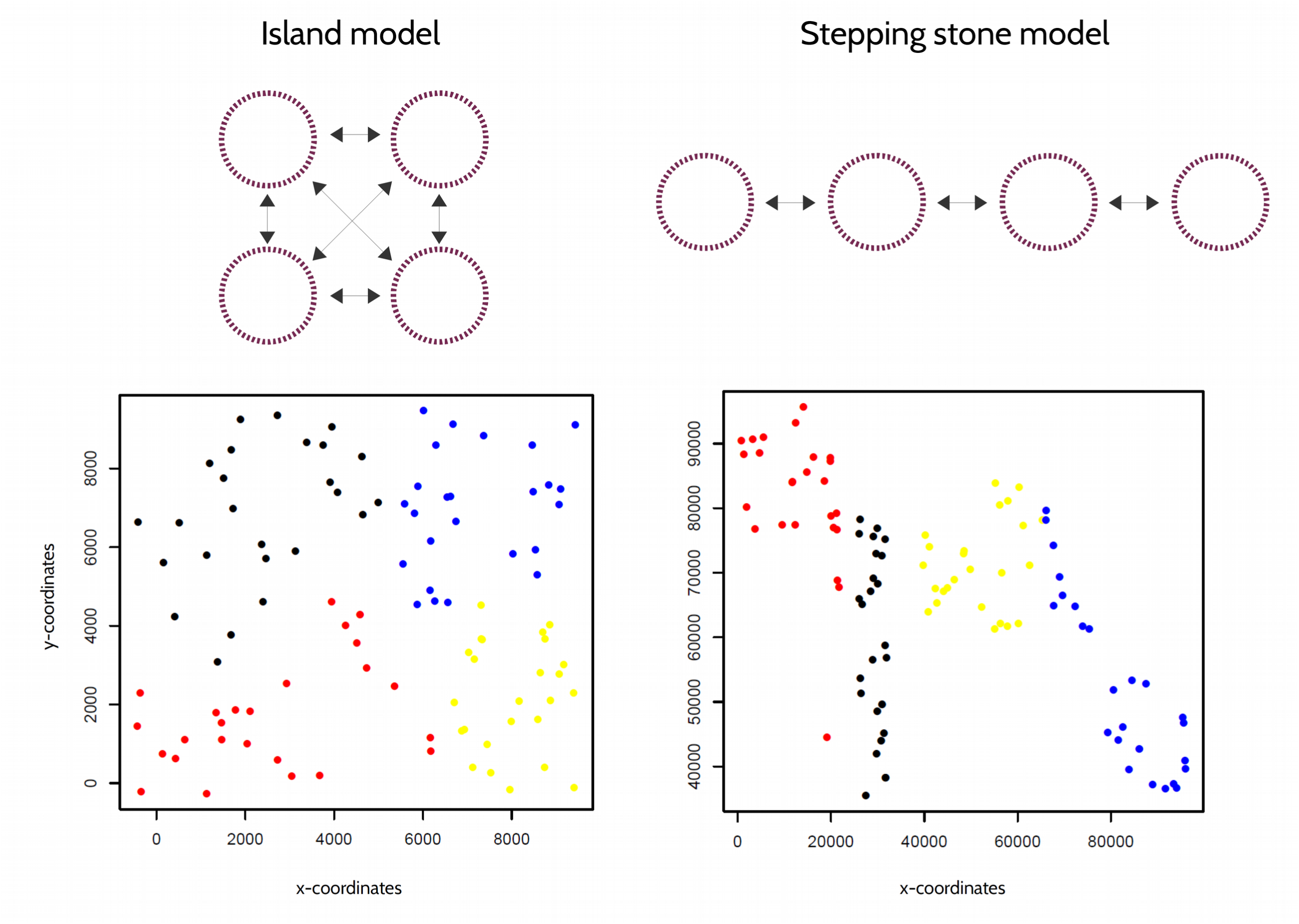
Graphical representation of island and stepping stone migration models (IS and SS) in the panel above. Black rows represent the presence and direction of migration rates among populations (purple circles). The panel below represents two examples of simulated global patterns, where a set of 100 pairs of coordinates are picked from a set of 1000 random pairs of coordinates built in 2D squares at different scales (in the example here reported the scales are 1:1e4 and 1:1e5, respectively). Every 25 pairs of coordinates are assigned to a different simulated population, distinguished by red, blue, black and yellow colors, in order to obtain spatially segregated populations. These simulated spatial distributions are used to calculate the matrix *L* of spatial connection (see Figure S1.

We tested 100 simulations each for all the 30 sets of geographic coordinates (random, positive and negative), for each of the three migration rates (0.005, 0.01 and 0.1), for each of the three migration models (IS, SS, IBD; total of 9,000 tests per migration model). We repeated all tests using a subset of 40 SNPs per individual, for a total of 18,000 tests in the absence of spatial structures, and and 36,000 tests in the presence of global or local structures.

## RESULTS

### Statistical power of the spca_randtest

We compared the performances of the *spca_randtest* with the *global* and *local* tests in three settings: in the absence of spatial structure, and in the presence of global, and local structures. The results obtained in the absence of spatial structure show that all tests have reliable type I errors (Table 1 and 2). The *spca_randtest* exhibited consistently better performances for detecting existing structures in the data than both *global* and *local tests* (Table 1 and 2). Although our simulated local spatial patterns turned out more difficult to detect than global patterns, the *spca_randtest* is twice to five times more effective than the *local test* (Table 1 and 2). Generally, the underlying migration model, the migration rate and the number of loci affect the ability of all tests to detect non-random spatial patterns. Both *spca_randtest* and *global* and *local tests* have in fact a lower sensitivity in presence of island migratory schemes, while results for stepping stone and isolation by distance models are more satisfying (Table 1 and 2). Increasing migration rates lead to a higher rates of false negatives for all tests, which can be overcome using more loci (Table 1 and 2).

**Table 1.**
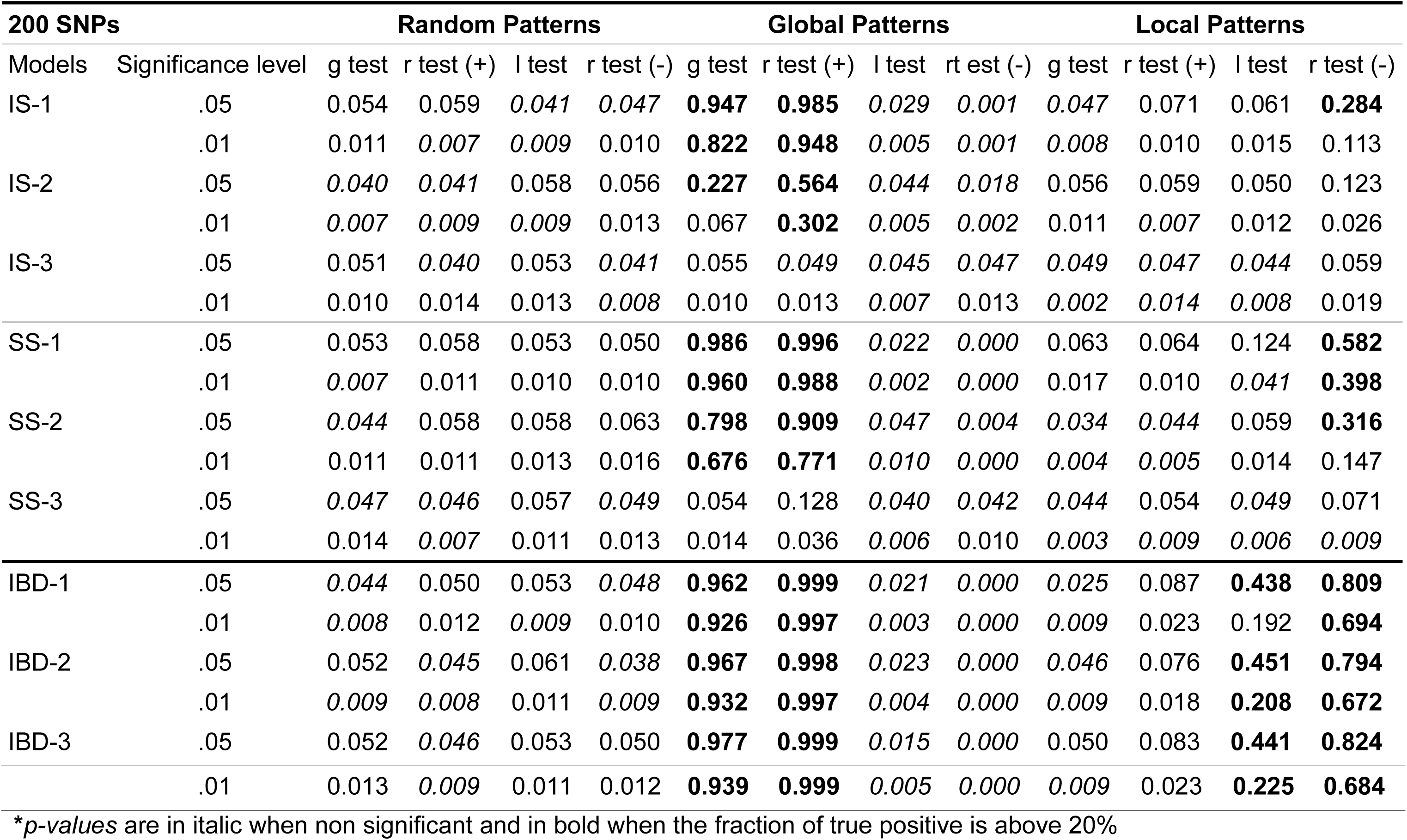
Significant results for g*lobal test* (g test), l*ocal tests* (ll test), and *spca_randtest* (r test +/-) for random, global and local patterns using 200 loci per individual. IS, SS, IBD indicate the migration models (see Methods); different migration rates are coded by number: 1 = 0.005, 2 = 0.01 and 3 = 0.1. Results show the proportion of significant tests over 1,000 replicates, based on 1,000 permutations with thresholds.05 and.01.

**Table 2.** Results for the same simulations reported in Table 1 using a subset of 40 loci per individual.

| 40 SNPs |  | Random Patterns |  |  |  | Global Patterns |  |  |  | Local Patterns |  |  |  |
| --- | --- | --- | --- | --- | --- | --- | --- | --- | --- | --- | --- | --- | --- |
| Models | Significance level | g test | r test (+) | l test | r test (-) | g test | r test (+) | l test | r test (-) | g test | r test (+) | l test | r test (-) |
| IS-1 | .05 | 0.052 | 0.061 | 0.046 | 0.050 | **0.591** | **0.807** | *0.033* | *0.004* | *0.036* | *0.000* | 0.055 | 0.077 |
|  | .01 | 0.016 | 0.013 | 0.010 | *0.007* | **0.393** | **0.592** | *0.005* | *0.000* | *0.004* | *0.000* | 0.015 | 0.022 |
| IS-2 | .05 | 0.053 | *0.047* | *0.038* | *0.042* | 0.103 | **0.226** | *0.046* | *0.020* | 0.073 | *0.000* | 0.057 | *0.038* |
|  | .01 | 0.011 | *0.009* | *0.006* | *0.006* | 0.022 | 0.072 | 0.011 | 0.005 | 0.012 | *0.000* | 0.010 | *0.006* |
| IS-3 | .05 | *0.047* | 0.050 | 0.050 | *0.045* | *0.048* | 0.060 | *0.044* | *0.042* | *0.036* | *0.000* | 0.053 | *0.026* |
|  | .01 | *0.009* | 0.011 | *0.008* | *0.007* | *0.009* | 0.011 | 0.011 | 0.011 | *0.002* | *0.000* | 0.013 | *0.001* |
| SS-1 | .05 | 0.052 | 0.054 | *0.039* | *0.049* | **0.898** | **0.949** | *0.017* | *0.000* | 0.050 | *0.001* | 0.067 | 0.169 |
|  | .01 | *0.009* | 0.012 | *0.005* | 0.011 | **0.826** | **0.865** | *0.006* | *0.000* | *0.007* | *0.000* | 0.021 | 0.052 |
| SS-2 | .05 | *0.046* | *0.045* | 0.050 | *0.046* | **0.528** | **0.588** | *0.044* | *0.009* | 0.052 | *0.000* | *0.048* | 0.081 |
|  | .01 | 0.013 | 0.010 | 0.010 | 0.015 | **0.377** | **0.370** | 0.016 | *0.000* | *0.005* | *0.000* | 0.011 | 0.014 |
| SS-3 | .05 | 0.068 | *0.040* | 0.050 | *0.048* | 0.066 | 0.055 | 0.053 | *0.033* | *0.026* | *0.000* | *0.047* | *0.023* |
|  | .01 | 0.014 | *0.005* | 0.013 | 0.012 | 0.012 | *0.009* | *0.005* | *0.006* | *0.006* | *0.000* | *0.008* | *0.000* |
| IBD-1 | .05 | *0.049* | 0.053 | 0.052 | 0.057 | **0.822** | **0.883** | *0.027* | *0.002* | *0.034* | 0.055 | 0.124 | **0.480** |
|  | .01 | *0.005* | *0.008* | 0.013 | 0.013 | **0.755** | **0.742** | *0.004* | *0.000* | *0.005* | *0.008* | 0.032 | **0.278** |
| IBD-2 | .05 | *0.043* | 0.054 | 0.060 | *0.049* | **0.835** | **0.880** | *0.028* | *0.001* | *0.043* | 0.051 | 0.111 | **0.458** |
|  | .01 | 0.011 | *0.007* | 0.015 | *0.009* | **0.755** | **0.732** | *0.005* | *0.000* | *0.008* | 0.015 | 0.026 | **0.259** |
| IBD-3 | .05 | *0.043* | *0.042* | 0.051 | 0.050 | **0.844** | **0.899** | *0.026* | *0.002* | *0.048* | 0.058 | 0.115 | **0.465** |
|  | .01 | 0.012 | 0.013 | 0.012 | 0.010 | **0.763** | **0.756** | *0.007* | *0.000* | *0.009* | 0.010 | 0.023 | **0.263** |

\**p-values* are in italic when non significant and in bold when the fraction of true positive is above 20%

Significant eigenvalues are assessed using a hierarchical Bonferroni correction which accounts for non-independence of eigenvalues and multiple testing (Figure 2). Strong patterns (e.g. IBD) tend to produce a higher number of significant components than weak patterns (e.g. island models with high migration rates), which are otherwise captured by fewer to no components.

### Application to real data

We have run the sPCA to compare the new *spca_randtest* and previous tests to a real dataset of human mitochondrial DNA (mtDNA). We used a dataset of 85 populations from Central-Western Africa that spans a big portion of the African continent (from Gabon to Senegal; Montano et al 2013). Previous analysis on these data detected a clear genetic structure from West to Central Africa with ongoing stepping stone migration movements. We therefore expected that this spatial distribution of genetic variation would be detected as significant. In the sPCA, populations were treated as units of the analysis, for which allele frequencies of mtDNA polymorphisms are calculated per population. The same approach was used in Montano et al 2013 to run a discriminant analysis of principal components (DAPC; Jombart et al 2010) and detect population genetic structure. The sPCA analysis is found non significant by *global* and *local* tests after 1e4 permutations (*p*-value > 0.5), while the *spca_randtest* detects a significant global pattern already with 500 permutations, and with 1e4 permutations the *p*-value for global patterns is 0.005. The second step of the test on single eigenvalues finds the three most positive components to be significant after Bonferroni correction (Table 3). Significant axes can thus be plotted against the spatial network to give a biological interpretation to the results (Figure 3).

**Table 3.**
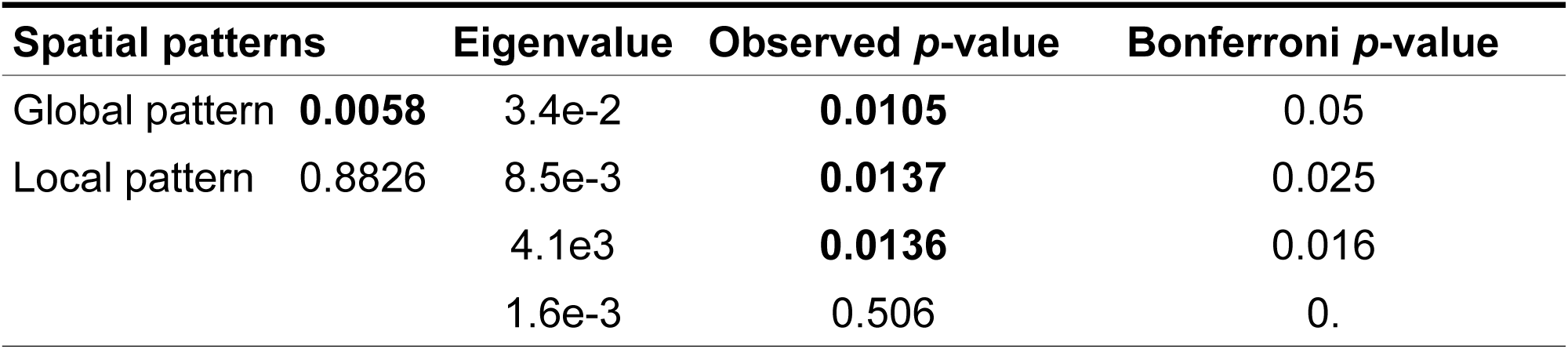
Results of the *spca_randtest* with 1e4 permutations on the human mtDNA dataset (Montano et al, 2013). The simulated distribution of the 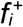 and 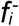 statistics are compared to the 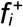 and 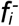 statistics observed for the original dataset. A significant global pattern (or significant 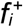 observed statistics) is found with the *spca_randtest* (p-value < 0.01). Thus, each eigenvalue is compared with its simulated distribution and assigned to be significant if its observed *p*-value is lower than the corrected Bonferroni *p*-value, with starting threshold of 0.05. Significant observed p-values as compared with Bonferroni corrected p-values are highlighted in bold.

**Figure 3.**
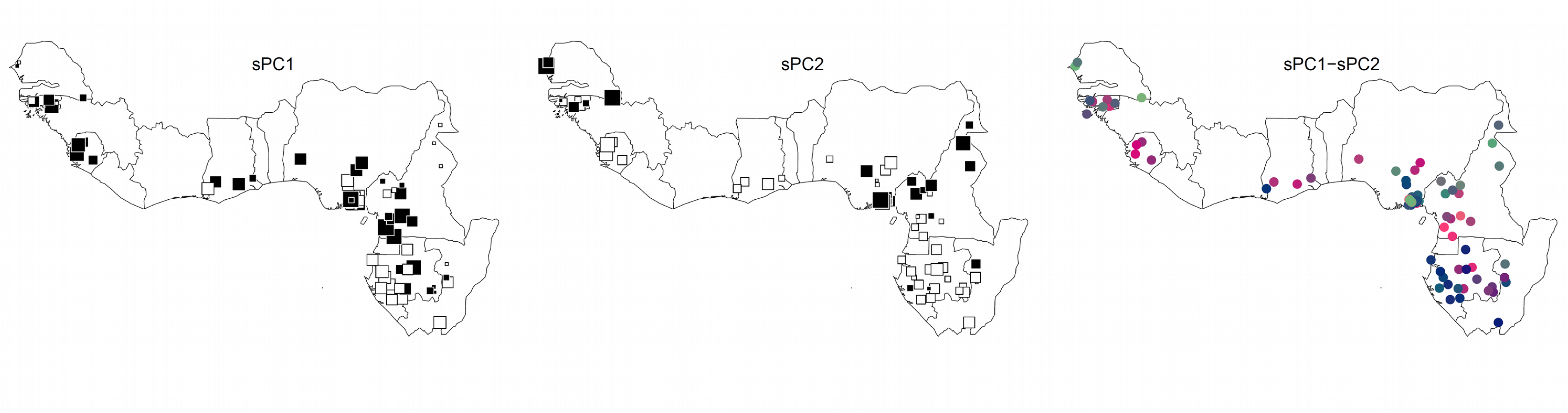
Plot of the first and second most positive observed eigenvalues of the mtDNA dataset here analysed. The background map represents the countries from where the populations included into the original study were sampled (from West to East: Senegal, Guinea-Bissau, Guinea, Sierra Leone, Liberia, Ivory Coast, Ghana, Togo, Benin, Nigeria, Cameroon, Equatorial Guinea, Gabon, Congo). sPC1 and sPC2 are represented independently using a square size proportional to the value of each population along the first and second component, respectively. Whites squares show negative values and black squares the positive values, with size being proportional to the absolute value of the coordinate. sPC1-sPC2 is a summarized representation of the values along the first and second component assumed by each population, using a color gradient.

## DISCUSSION

We introduced a new statistical test associated to the sPCA to evaluate the statistical significance of global and local spatial patterns. Using simulated data, we show that this new approach outperforms previously implemented tests, having greater statistical power (lower type II errors) whilst retaining consistent type I errors. Our simulations also suggest that demographic settings and migratory models can substantially impact the ability to detect spatial patterns. Indeed, high migration rates, non-hierarchical migration models, such as island model, and low amount of loci can hamper or worsen the performance of the test, preventing the detection of actual spatial patterns. In lack of previous information on the demographic history and/or the movement ecology of the population under study, it is certainly useful to exploit all the available genetic information. In this regards, our simulations show how an increased number of loci does improve the ability of the test to provide meaningful results.

The impact of specific factors such as the effective population size or the number of individuals sampled per population remain to be investigated. A more extensive simulation study, possibly comparing different non-model based methods such as sPCA, would clarify the extent of the spatial information that can be obtained with such methods without comparing explicit evolutionary hypotheses. In fact, the sPCA and the associated *spca_randtest* cannot distinguish between explicit migration models. However, the possibility to detect which eigenvalues contain the spatial information provides the user with further information to interpret the biological meaning of the spatial structure, by focusing on few meaningful dimensions.

Our data application seems to confirm that the *spca_randtest* is more effective than *global* or *local* tests. We chose indeed a previously published dataset of human populations which span a subcontinental area of Africa and had been originally detected to be a highly structured dataset with a geographic cline of population differentiation (Montano et al 2013). On the basis of the original results, we would have expected a spatial global structure to be present in the data and thus detected with an sPCA. While the *global* test failed to provide statistical significance, the *spca_randtest* did obtain significant results and pointed to the three first most positive components to be also significant after Bonferroni correction. In agreement with the original interpretation of the genetic structure within the samples, spatial component 1 (SP1) shows a clear differentiation of populations in the Gabon-Congo region, while SP2 detects differentiation of Central Nigerian and North Cameroonian populations, on one hand, and extreme Western populations of Senegal, on the other hand (Figure 3). The colored combination of the first and second most positive component (Figure 3) also correctly detects a more fragmented differentiation across Central forested areas (Cameroon, Gabon and Congo) compared to more homogeneous Central-Western populations, which was the main result of the original publication based on very different approaches (Montano et al 2013). We limited the analysis to these two component as the third did not add much information to the previous.

Our simulation approach coupled with a real data application well illustrates the informativeness of our new test to retrieve significant spatial patterns, being these global or local structures and highlights the usefulness of selecting a specific number of significant components to interpret the biological meaning of the results.

## Declarations

### Data Accessibility

https://github.com/thibautjombart/adegenet/blob/master/R/spca_randtest.R

### Acknowledgements

The authors declare no conflict of interest

### Author contributions

Test development: VM and TJ. Data analysis: VM. Wrote the manuscript: VM and TJ.

---

**Figure S1.**
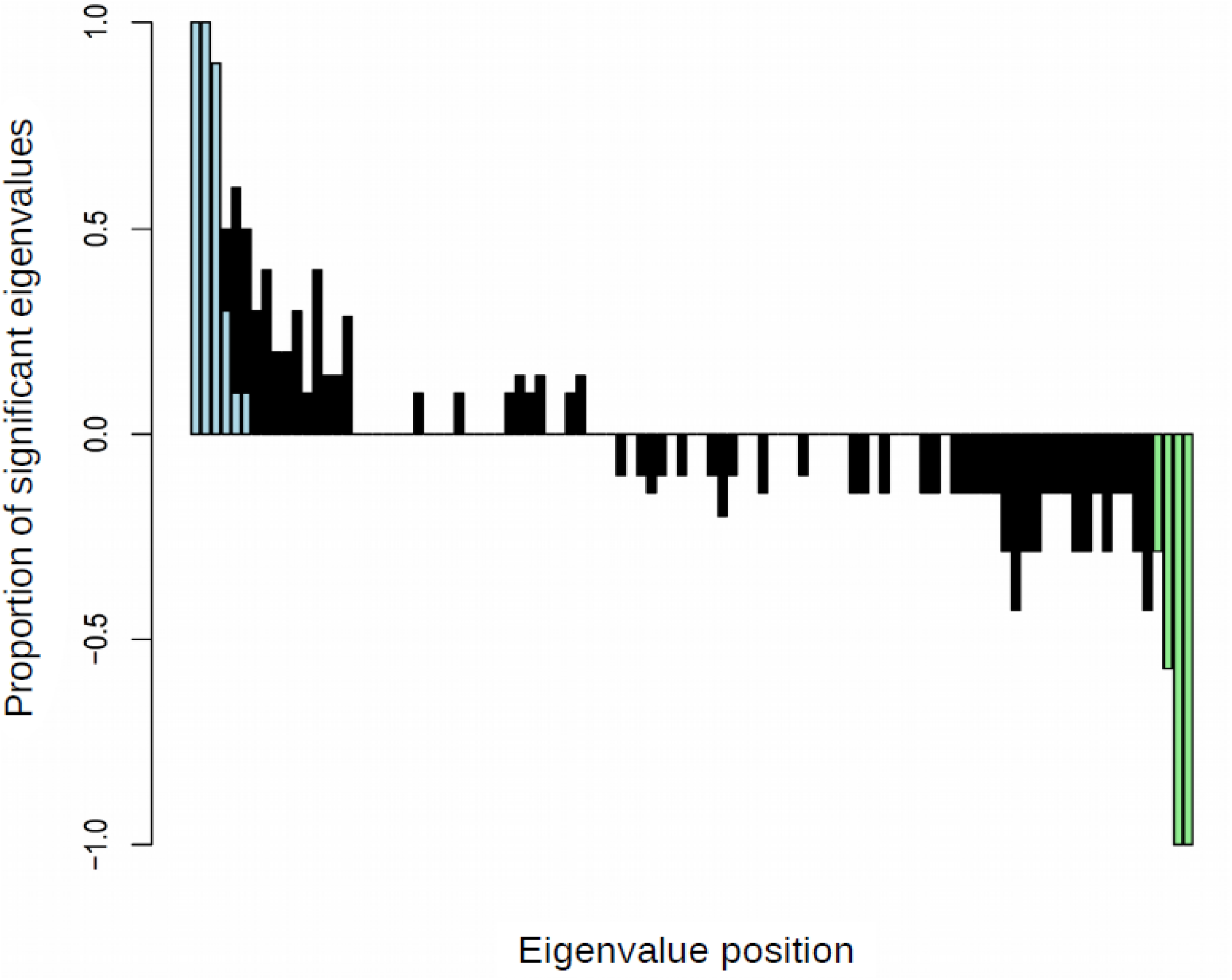
Distributions of significant eigenvalues detected in the presence of global (blue bars) and local (green bars) spatial patterns after hierarchical Bonferroni correction, for 100 significantly positive and 100 significantly negative patterns. Black bars correspond to eigenvalues which are significant without Bonferroni correction. Bars’ height indicates the frequency of observing a significant eigenvalue in a certain position (from most positive to most negative) over the 100 tested patterns.

